# Juvenile hosts and natal dispersers are protected in the early stages of epidemics

**DOI:** 10.64898/2026.02.04.703830

**Authors:** Louis T. Bubrig, Irish A. Amundson, Sarah K. Talley, Sophia G. Kuzminski, Amanda K. Gibson

## Abstract

Parasite prevalence varies in time and space. Thus, hosts may escape infection by dispersing out of habitats where parasites are present. However, it is not clear if the advantage of avoiding parasites outweighs the cost of dispersing. Juvenile hosts are expected to be relatively protected from environmentally-transmitted parasites, and we hypothesize that this age bias in transmission could magnify the benefits of juvenile (*i.e.,* natal) dispersal. We tested these ideas in the model nematode *Caenorhabditis elegans*, a host with discrete life stages and natal dispersal, and its environmentally transmitted microsporidian parasite *Nematocida parisii*. We found that under standardized exposure conditions, larger *C. elegans* individuals (corresponding to older life stages) acquired many more parasites than smaller (younger) individuals. We found this same bias during multigeneration epidemics, especially during early stages of the epidemics. We also found that *C. elegans* dispersal larvae were less likely to be infected and harbored less severe infections than the population mean. We conclude that the early stages of an epidemic can provide young hosts with a window of opportunity to escape infection by dispersing.

## Introduction

Dispersal is the movement of individuals between habitats (Bowler and Benton 2005) and is a critical component of life history (Denno et al. 1991). Theory proposes that dispersal evolves so hosts can escape habitats that degrade over time (Van Valen 1971). In many species, dispersal is performed by juveniles (“natal dispersal”), which allows them to escape poor conditions and reproduce elsewhere (Clobert et al. 2001). A challenge in studying dispersal is identifying the specific drivers of habitat decline that could confer an advantage to individuals that leave (Friedenberg 2003; Starrfelt and Kokko 2012).

Parasites are one prominent possibility (Connell 1971; Augspurger 1983; Boulinier et al. 2001; van Baalen and Hochberg 2001). Parasites impose costs on their hosts, and the risk of infection increases rapidly during an epidemic (Hochberg et al. 1992; Agnew et al. 2000). Therefore, hosts that disperse during an epidemic could benefit by avoiding contact with their parasite (Curtis 2014). Ultimately, dispersal is evolutionarily favored only if this benefit of escaping infection outweighs the benefit of remaining in the habitat to reproduce (Boulinier et al. 2001). This balance depends on the epidemiological context: which hosts are currently infected, which will be infected next, and how rapidly is the disease spreading? By this argument, epidemiological dynamics are key to understanding dispersal as a parasite escape strategy.

A key feature of epidemics is biased transmission among host groups. Host age influences transmission (Grenfell and Anderson 1985; VanderWaal and Ezenwa 2016; McCallum et al. 2017). Disease burden is predicted to increase with age when hosts acquire parasites from the environment: older life stages are often larger and thus are more exploratory and present a larger surface area for parasites to contact (Mohr 1961; Randolph 1975; Boyer et al. 2010; VanderWaal and Ezenwa 2016; Shaw et al. 2024). Body size and disease burden can exhibit a superlinear relationship (Mohr 1961; McCallum et al. 2017; Shaw et al. 2024), meaning that even host species with modest growth from juvenile and adulthood can experience a pronounced transmission bias. Additionally, older hosts have a longer cumulative period of exposure to parasites than younger hosts. Thus, we expect reduced disease burden in juveniles compared to adults because 1) they acquire parasites at a lower rate and 2) their cumulative exposure to the parasite is shorter. In species with natal dispersal, juveniles disperse. Therefore, we expect the same transmission biases that shield juveniles from infection within a patch could also protect dispersers leaving that patch, deepening the advantage of dispersal.

To investigate the intersection between transmission biases and dispersal, we tested two hypotheses. First, we hypothesized that smaller, younger hosts should be less burdened with environmentally-transmitted parasites than larger, older hosts. Second, we hypothesized that natal dispersers should be less burdened with parasites compared to the populations they left. We tested these hypotheses in controlled exposure assays and in experimental epidemics using the nematode *Caenorhabditis elegans* and its environmentally-transmitted natural parasite *Nematocida parisii*. The host, *C. elegans*, disperses in nature using a specialized larval stage (*i.e.,* the *dauer*) that precedes reproductive adulthood. We used two strains (genotypes) of *C. elegans* that vary in resistance to *N. parisii* to assess the generality of our results. We found that juvenile hosts faced a lower risk of infection in the early phases of epidemics. Additionally, we found that dispersers were less likely to be infected and harbored smaller infections than the populations they came from.

## Materials and Methods

We first tested how parasite contact rate differed among host life stages in a controlled spore acquisition assay. Then, we evaluated infection prevalence and severity among host sizes/ages over the course of epidemics. Finally, we extracted dispersal larvae from host populations experiencing these epidemics to measure infection prevalence and severity in natal dispersers.

### Natural History

*C. elegans* progresses through four larval stages (named L1-L4) and one adult stage. Host length varies along a continuous spectrum. Hosts molt into the next larval stage upon reaching predictable lengths (Byerly et al. 1976). *C. elegans* also has a specialized dispersal stage called the dauer. In nature, small numbers of dauers colonize patches of bacteria-rich rotting vegetation. Many patches are colonized by individuals of a single selfing lineage (Richaud et al. 2018; Sloat et al. 2022) which establish rapidly growing, mixed-stage populations (Frézal and Félix 2015). Increasing population density and decreasing food trigger juveniles (specifically the first two larval stages, L1 and L2) to develop into dauer larvae, which engage in natal dispersal (Cassada and Russell 1975). Because *C. elegans* inhabits ephemeral resource patches, dispersal is critical to the persistence of lineages in metapopulations.

*C. elegans* is commonly infected by the microsporidian *Nematocida parisii*. Nematodes ingest *N. parisii* spores while feeding. *N. parisii* infects the intestine and, after about two days, sheds tens of thousands of spores back into the environment (Troemel et al. 2008; Szumowski et al. 2012). Due to *C. elegans’* rapid life cycle (see Table S1), epidemics observed in the lab span multiple host generations and therefore coincide with demographic changes and population expansion. *N. parisii* can infect all nondauer life stages, and hosts infected in pre-dauer stages (*i.e.*, L1 or L2) retain the infection when developing into dauers. Infected dauers can transmit infection to new colonies (*personal observation*). However, dauers do not feed (Cassada and Russell 1975) and therefore cannot acquire infection if they were not previously infected as juveniles.

### Experimental Set-up: Spore Acquisition Assay

We conducted all experiments with the standard *C. elegans* lab strain N2, which suffers relatively large costs when infected with *N. parisii*. For the epidemic experiment, we also included the strain CB4856, which is considered more resistant to *N. parisii* (Balla et al. 2015; Bubrig et al. 2022). We thawed nematodes from −80°C stocks for each experimental block. After thawing, we propagated populations at 20°C and transferred lines to fresh solid media (100-mm Nematode Growth Media (NGM Lite, US Biological) plates seeded with 300 µL *Escherichia coli* OP50 food) every 3-4 days for approximately three weeks. We generated *N. parisii* spore stocks by following a previously reported protocol (Bubrig et al. 2022). The spore acquisition assay used varying volumes of a spore stock with a concentration of 108,833.21 spores/µL. The assay’s control treatment used control lysate, which was made similarly to spore stocks except using uninfected host populations. The epidemic experiment used a spore stock with a concentration of 260,607.42 spores/µL to deliver an initial dose of 100 spores/mm^2^ of plate area, which is considered a very low dose (Willis et al. 2021).

To test whether different host life stages acquired parasite spores at different rates, we exposed three life stages (L1, L4, and adult) to parasites for a fixed exposure window. Life stages were acquired by extracting eggs with a standard bleach protocol (Stiernagle 2006) and allowing hosts to develop for different intervals of time, corresponding to the developmental timings in Table S1. Within each block, two populations of L1s, two of L4s, and two of adults were suspended in M9 buffer. We combined 1200 worms from each population with 150 µL 1x *E. coli* OP50 superfood diluted in S-medium, and the relevant exposure treatment onto 60-mm NGM plates. Control hosts were treated with 40 µL control lysate, and exposed hosts were treated with 20µL, 40µL, or 80µL of spore stock for 3 hours at 20°C before being washed into M9 and preserved in acetone. Samples were kept at −80°C until fluorescence *in situ* hybridization (FISH; see *Microscopy and FISH* below) to determine the number of spores acquired. Across three experimental blocks, L1 groups were replicated four times, L4s at least four times, and adults three times.

### Experimental Set-up: Epidemic Experiment

To determine whether disease burden in epidemic contexts was biased by host size/age, we established host populations with *N. parisii* epidemics. Our design replicated how we expect *C. elegans* and *N. parisii* interact in the wild. We isolated host eggs as above and plated them for 48 hours, then transferred 20 L4 individuals to each experimental plate (n = 39 plates for each strain). Three days after this transfer, we introduced 3.01 µL of spore stock to each plate, spread it evenly, and allowed plates to dry. This three-day delay ensured that at the start of the epidemic, the host population was actively proliferating with a variety of life stages present (Figure S1). We report time counting from the day populations were exposed to spores (*e.g.,* day 1 of the epidemic is the day after spores were introduced).

Data were collected on days 1 through 5 of the epidemic, plus days 8 and 10. Each host strain was tested in two blocks, but day 10 in N2 and day 1 in CB4856 appear in only one block. At each timepoint, three experimental plates were chosen at random. We washed each population into 5 mL M9, then diluted with more M9 until a ∼20 µL aliquot would contain a few dozen individuals. To estimate population size, we counted all live hosts in 4 of these small aliquots. Then, we removed a sample of the population and stored it in acetone in a −20°C freezer for FISH. We then extracted dauers from the remaining population by treating the populations with 1% SDS for 15 minutes to kill nondauer life stages (Cassada and Russell 1975). We halted SDS treatment by washing the population with M9 five times. The pellet was spotted onto unseeded 100-mm NGM plates, dried, and incubated for 1 hour at 20°C to allow live dauers to crawl from the pellet. We cut out the pellet from the plate, then washed live dauers into M9 and quantified them in four small aliquots per population. Finally, we removed a sample of the dauers and stored them in acetone at −20°C for FISH. The sample taken before the SDS wash is called the “whole population” and the sample taken after is called “dauer.”

### Microscopy and FISH

Our goal was to investigate the relationship between host size/age and infection. We evaluated both factors via microscopy. We processed acetone-stored samples with FISH probes which hybridize to *N. parisii* rRNA and reveal active *N. parisii* infections (Troemel et al. 2008). Hosts were deposited onto thin 3% agarose pad microscopy slides and imaged under a fluorescent source with a Texas Red filter (Microscope: Leica DM6 B; Camera: Leica DFC3000 G).

For the spore acquisition experiment, at least 12 hosts per sample were scored by manually counting individual fluorescent spores. For the epidemic experiment, we imaged hosts and used ImageJ to measure their length from nose to tail. These lengths were used in statistical analyses. However, for figures we estimated each host’s life stage (L1-L4 or adult) using characteristic molting lengths (Table S1). Note that dauers in whole population samples were classified as either an L2 or L3 due to dauers’ sizes overlapping with those stages’ size ranges. Next, we calculated infection load (percent body area infected) by adding up the total area of all fluorescent patches inside a host and dividing by the host’s body area. Finally, we measured fluorescence intensity specifically along the host’s head-to-tail axis and calculated “linear load,” or the percentage of this axis that contained infection. Linear load is an alternative load metric that should be robust to dauers radially constricting their gut, a process which may artificially deflate the area-based load metric. We scored 5,322 individual nematodes from 78 populations.

### Statistical Analysis

Data analysis and modeling were performed in R version 4.4.0 using the glmmTMB package (Brooks et al. 2017). For all datasets, we accounted for variability among laboratory plates by modeling plate ID as a random effect. We modeled spore acquisition data using negative binomial distributions (Table S2), prevalence using binomial distributions (Tables S2, S4), and load using beta distributions (Tables S3, S5, S6). For all datasets, we built an unrestricted model with our variable of interest (*e.g.,* Host Length) and other covariates. We tested each unrestricted model for zero inflation and over- or under-dispersion using the DHARMa package (Hartig 2016) and corrected deviations using zero inflation and dispersion formulas (Tables S3, S5, S6). In analyses comparing whole populations to dauers, we included an SDS-by-Strain interaction since the health of dauers appeared to depend on strain. We tested whether interactions were significant by performing an ANOVA between models with and without the interaction. If the interaction was supported, we split the dataset into an N2 dataset and a CB4856 dataset, then tested whether the main effect of SDS was supported for each strain separately. In these cases, we accounted for multiple comparisons using Bonferroni correction to multiply *p*-values by the number of ANOVA tests performed (*e.g.* by 3 in the case of testing for an interaction, then testing the main effect of SDS in two strains).

## Results

We tested two interrelated hypotheses regarding size/age bias in disease spread. First, we investigated whether small/young hosts were relatively less burdened by parasites in a controlled infection assay and an experimental epidemic. Second, we examined dispersers’ infection status to determine if they were relatively less burdened than their populations of origin.

### Differences in spore acquisition by life stage

We quantified parasite spores acquired by hosts of three life stages during a brief 3-hour exposure period. Spore acquisition differed among life stages (Life Stage: *χ*^2^ = 62.7, *df* = 2, Bonferroni-corrected *p <* 0.001; Table S2). L1 hosts acquired fewer spores than later life stages (Figure 1). On average, L4s acquired 3.6 times and adults 7.2 times more spores than L1s (Table S2). At the highest dose, L1s acquired a maximum of 23 spores, compared to 205 for L4s and 227 for adults.

**Figure 1.**
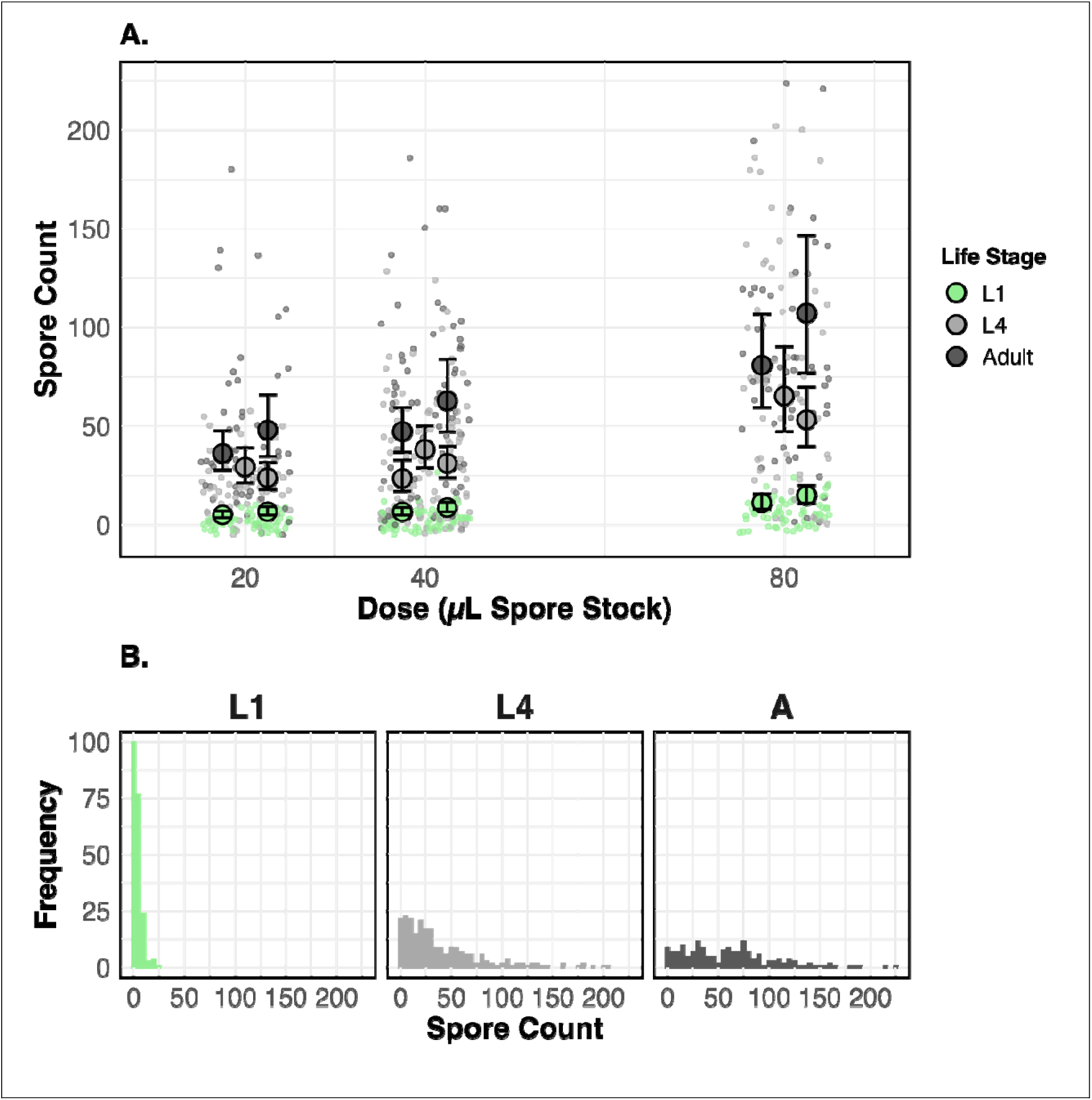
Spore acquisition in *C. elegans* life stages. *A*: Number of *N. parisii* spores acquired by L1, L4, and adult hosts after a 3-hour exposure period at three parasite doses. Small shaded points show data for individual hosts. Larger points show mean spore acquisition within an experimental block, and error bars show 95% confidence intervals. *B*: Histograms showing the distribution of spores acquired by individual hosts across the three tested life stages, pooled across doses.

### Does host size/age influence the likelihood and severity of infection?

We first described general epidemic dynamics in the *C. elegans*-*N. parisii* system without regard to host size/age. Infection prevalence increased over time, with each additional day increasing the odds of being infected by 2.5 (Table S3) and infection load by a factor of 1.3 (Table S4). However, prevalence did not increase monotonically. In N2, prevalence rose to 30.4% ± 7.3% on day 1 of the epidemic, dropped to 5.3% ± 1.9% on day 2, then eventually plateaued around 80-90% for the remainder of the timeseries. Similarly, prevalence in CB4856 rose initially, declined to 15% for two days, then increased to the 50-70% range (Figure 2). N2’s drop in prevalence on day 2 and CB4856’s drop on days 2 and 3 coincided with rapid population growth (Figure S2).

**Figure 2.**
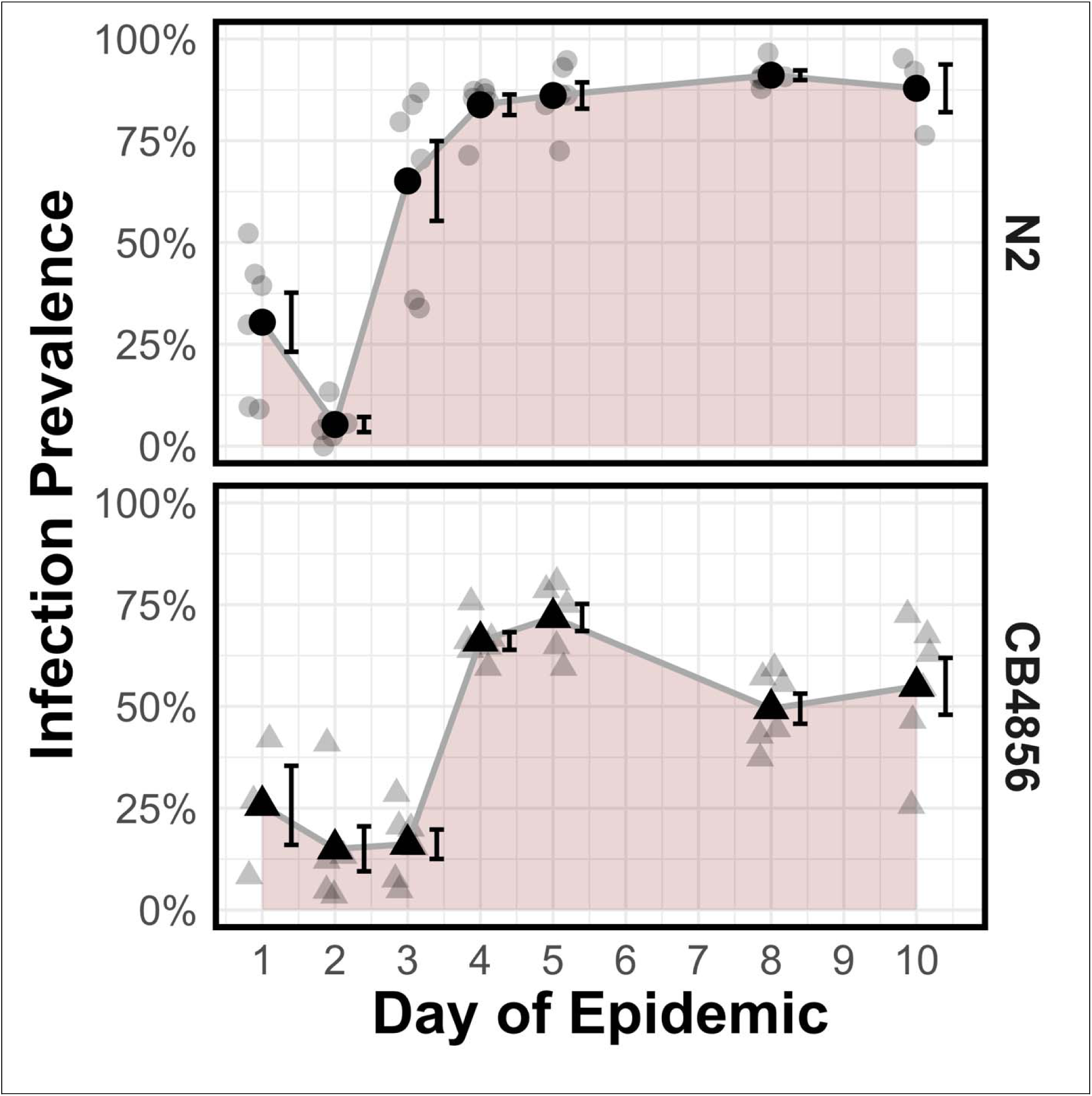
Infection prevalence during epidemics. Prevalence is calculated as the percent of hosts that are infected. Faded grey points show the prevalence for individual replicate plates. Black points show the prevalence averaged across replicate plates (n = 6 per time point, except n = 3 for N2 day 10 and CB4856 day 1). Error bars show standard error of the mean.

When then measured host size to determine whether infection was biased among hosts of different sizes. Smaller body sizes correspond to more juvenile hosts, while larger body sizes correspond to later life stages (see *Materials and Methods*; Figure S1). During the epidemic, host populations grew rapidly (Figure S2) and transitioned from harboring all life stages to being juvenile-dominated during population expansion (Figure S1), which matched *a priori* expectations. Dauers started to form on day 3 in N2 and day 4 in CB4856 (Figure S3). We found that early in the epidemic, larger hosts (corresponding to later life stages) were more likely to be infected than smaller (juvenile) hosts. Using hosts’ lengths to estimate their life stage, we found that in N2 populations, no L1s or L2s were infected (prevalence = 0%) on day 1 of the epidemic, while 48.2% ± 10.1% of adults were. The difference was more pronounced on day 2: between 0% and 0.5% of L1s, L2s, and L3s were infected compared to 79.2% ± 16.4% of adults (Figure 3). Our statistical model estimated that every 100 µm increase in length increased a host’s odds of being infected by 2.3 (Table S3). Among the infected hosts, larger hosts had more severe infections: every 100 µm increase in length increased hosts’ parasite load 1.1-fold (Table S4; Figure S4). CB4856 showed a similar pattern to N2, with the smallest hosts (corresponding to L1s) retaining relatively low prevalence through the first five days of the epidemic. Infection prevalence became more uniform across host size as epidemics progressed (Figure 3), as suggested by a negative interaction between time and host length (DayOfEpidemic* RescaledLength: *χ*^2^ = 63.4, *df* = 1, *p <* 0.001; Table S3). In contrast, the relationship between host length and parasite load was robust to the progression of the epidemic (DayOfEpidemic* RescaledLength: *χ*^2^ = 3.3, *df* = 1, Bonferroni-corrected *p =* 0.135; Table S4).

**Figure 3.**
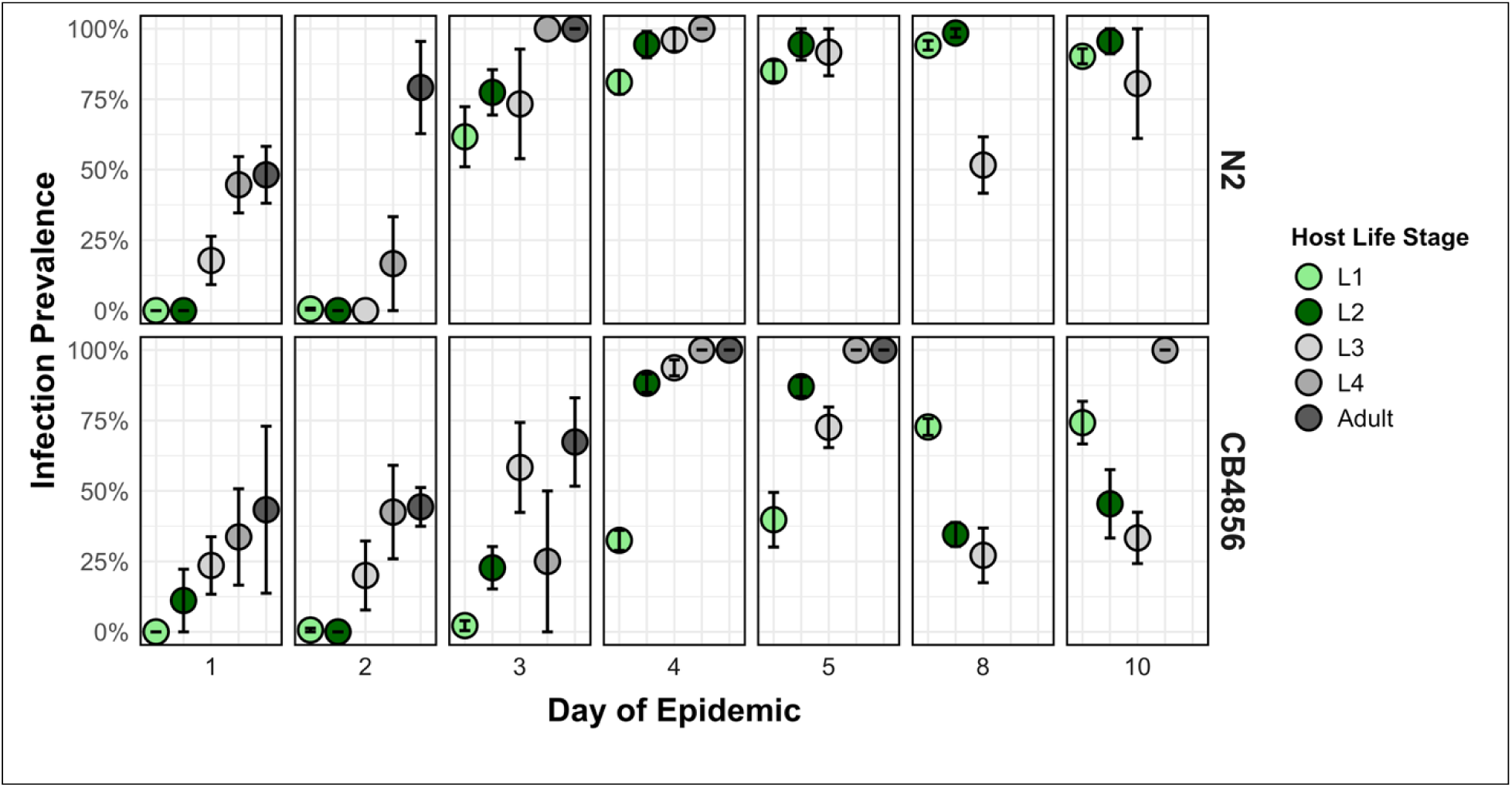
The infection prevalence of each life stage in the populations over time. Each panel shows data from a single time point. Each host’s length from nose to tail was used to estimate its life stage based on characteristic molting lengths. Datapoints that are missing for particular life stages indicate that no members of that life stage were found at that time point. Error bars show standard error of the mean across replicate plates. Missing error bars indicate that the age class was only found on one of the replicate plates.

### Are dispersers healthier than the general population?

To determine if dispersers were healthier than the population average, we extracted dauers and quantified their infection prevalence and load. On the first day of dauers in N2, day 3, only 18.8% ± 5.4% of dauers are infected while 65.1% ± 9.8% of individuals in the whole population are. Prevalence in dauers then increased and plateaued around 75%, around 10% lower than that of the whole population. In CB4856, prevalence in dauers began at a higher value than N2 (50.9% ± 6.7%) but fell to around 30% later in the epidemic and was consistently 10-20% lower than that of the whole population (Figure 4). When considering each replicate population and its dauer pool individually, most replicate populations followed this general pattern (Figure S5). Prevalence differed between dauers and the populations they came from (SDS: *χ*^2^ = 143.4, *df* = 1, *p <* 0.001) with individuals in the whole population having 2.5-fold higher odds of being infected than dauers (Table S5).

**Figure 4.**
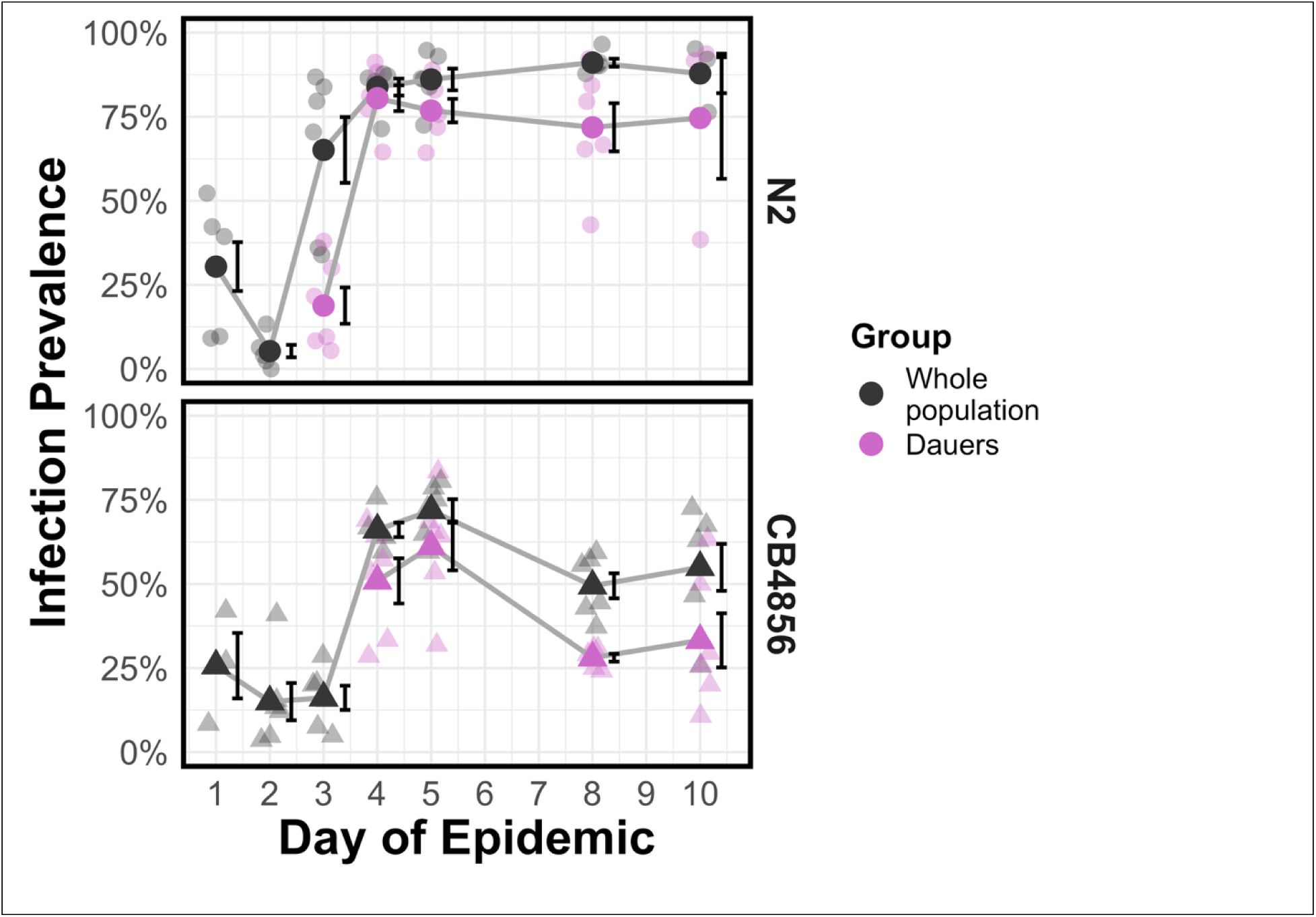
Infection prevalence of dispersing hosts relative to the population. Prevalence was calculated as the percent of hosts that are infected. Whole population data (black) may contain any *C. elegans* age classes including normal lifecycle larvae and dauers. Data for dispersers (pink) contain only individuals who survived SDS treatment, *i.e.* dauers. Days with no dauers in the population (see Figure S3) have no prevalence data. Faded points show data for individual replicate populations. Solid points show the mean across all replicate plates. Error bars show standard error of the mean.

We then tested the parasite load of infected dauers versus the populations they came from (Figure 5). Load was lower in infected dauers compared to the whole population. The magnitude of this difference depended on host strain (Strain*SDS: *χ*^2^ = 21.6, *df* = 1, Bonferroni-corrected *p <* 0.001; Table S6). Parasite load in early dauers (day 3 in N2, day 4 in CB4856) was low, with less than 1% of the host’s body area infected. In N2, load in dauers reached a maximum of 9.4% ± 6.3% compared to a maximum of 18.3% ± 3.8% in the whole population. In CB4856, load in the whole population reached a maximum of 5.5% ± 3.5% but never progressed higher than 1.1% ± 0.3% in dauers. Even in the latest time point, CB4856 dauers typically have small, isolated infections while N2 dauers have widespread infections throughout the gut (Figure S6). In modeling, the load of infected CB4856 dauers was lower than of the whole population by a factor of 2.8 (compared to a factor of 1.4 in N2), suggesting that CB4856 dauers have especially small infections compared to N2 dauers (SDS: *χ*^2^ = 114.5, *df* = 1, Bonferroni-corrected *p <* 0.001; Table S6). These results generally held when considering each population and its dauer pool individually (Figure S7). We also tested whether dauer loads were lower simply because dauer guts become radially constricted compared to other life stages. We calculated a “linear load” metric (see *Materials and Methods*) which quantifies the percentage of the host’s long axis that contains infections. We again found that dauers had less severe infections than the whole population (Table S7), and especially so in CB4856 (Figure S8).

**Figure 5.**
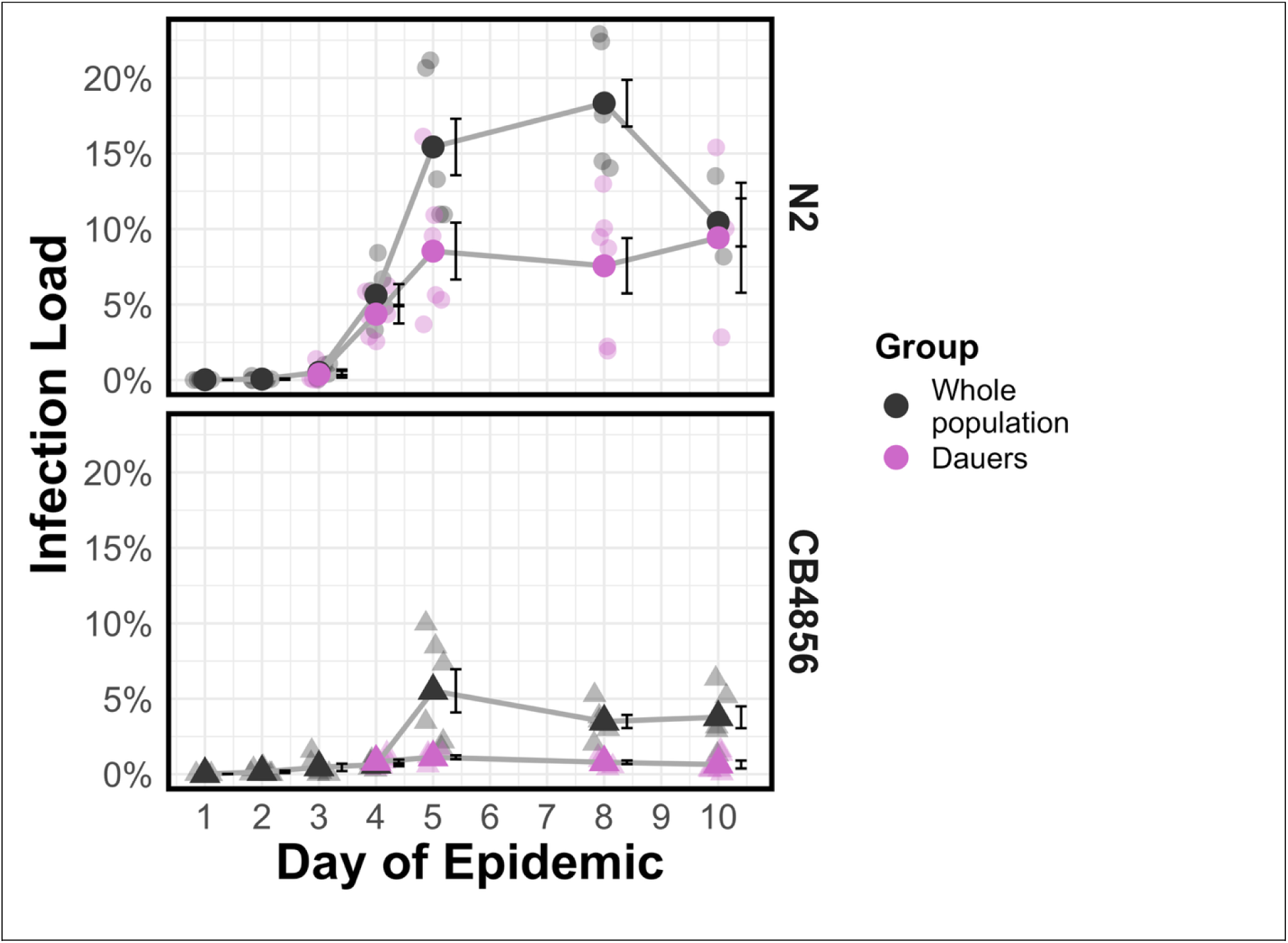
Parasite load in the whole population versus the dauer pool. Parasite load was estimated by dividing the area of fluorescence inside a nematode by the nematode’s body area. Whole population data (black) may contain any *C. elegans* age classes including normal lifecycle larvae and dauers. Dauer pool data (pink) contain only individuals who survived SDS treatment, *i.e.* dauers. Days with no dauers in the population (see Figure S3) have no load data. Faded points show data from a single replicate population. Solid points show the mean across all replicate plates. Error bars show standard error of the mean.

## Discussion

We used the tractable model host system *Caenorhabditis elegans* and its parasite *Nematocida parisii* to test for size/age biases in disease spread and its effects on dispersal in controlled short-term assays and long-term epidemic studies. We tested two hypotheses. First, that small/young hosts should be relatively less burdened with environmentally-transmitted parasites than larger/older hosts. Second, that juveniles who go on to disperse should be healthier than the population average. We found support for both hypotheses, suggesting that size and age bias in parasite transmission can deepen the advantage of dispersing during an epidemic.

We tested the first hypothesis, that smaller/younger hosts are less burdened by parasites than larger/older hosts, with a parasite acquisition assay. We found that older host groups acquired substantially more parasite spores than juvenile hosts (Figure 1). The standardized 3-hour exposure window meant that all host groups had the same cumulative exposure. Therefore, we inferred that older hosts contacted substantially more spores. We anticipate that this relationship is primarily allometric: the tight correspondence between age and size in *C. elegans* meant that individuals in older host groups were physically much larger than those in younger host groups. Previous work connects the rate of parasite contact to the surface area of the host’s contact surface. For example, larger rats acquire more chiggers in part because of their larger ventral area brushing against chigger-laden vegetation (Mohr 1961). Similarly, the host snail *Biomphalaria glabrata*’s contact with skin-burrowing schistosomes increases with its external surface area (Shaw et al. 2024). We expect that our spore acquisition results are analogous: *N. parisii* contacts *C. elegans* in the gut (Troemel et al. 2008), which increases in area and volume as hosts progress to adulthood. The relatively low throughput of the juvenile gut could explain why even at the highest parasite dose L1s contact few *N. parisii* spores (Figure 1). Age bias in parasite load is often intertwined with differences in immune function among different age groups (Grenfell and Anderson 1985). However, we think immune factors are unlikely to explain the differences we observe in this study. The exposure window was long enough for *N. parisii* to establish in host gut cells but too short to activate known host immune pathways that could fight infection (Balla et al. 2015).

We then set up replicate epidemics to test whether this juvenile protection held in a more dynamic exposure regime. The early stages of the epidemic occurred alongside host population growth (Figure S2). Theory has shown that parasite prevalence can temporarily decrease or stall early in epidemics when the host population grows faster than parasites can transmit (Bubrig and Gibson 2025). Our experimental populations exhibited this phenomenon, as rapid population growth appeared to temporarily suppress parasite prevalence in both host strains. The effect was more pronounced in N2 (Figure 2, S2), likely driven by the strain’s faster growth (Zhang et al. 2021). Prevalence then rapidly increased and plateaued. Consistent with the host strains’ reported resistance phenotypes (Balla et al. 2015), epidemics in the more susceptible N2 strain reached higher levels of prevalence than those in the more resistant CB4856 strain (Figure 2). As in the spore acquisition assay, smaller hosts (corresponding to juveniles) had lower odds of being infected than larger hosts during an epidemic. Of the hosts that were infected, smaller hosts had smaller infection loads than larger hosts.

The epidemic experiment added several new factors that were absent in the spore acquisition assay. The exposure period was prolonged from three hours in the assay to ten days, covering multiple host generations and allowing for host immunity to act. L1 larvae have the unique ability to clear *N. parisii* infections, but this cannot fully explain the relative health of juveniles observed in both strains (Figure 3), since this clearance ability is only seen in the CB4856 host strain (Balla et al. 2015). Additionally, we found that the magnitude of size bias in parasite prevalence was strongest in the early days of the epidemic and became less pronounced as the epidemic progressed. Epidemics had an early “grace period” (two days in N2, three days in CB4856) in which parasites transmitted among older hosts while largely sparing juveniles.

After this grace period, however, size bias in parasite prevalence collapsed, which may be explained by the accumulation of spores over time. While dose in the spore acquisition assay was constant, the dose experienced by hosts in the epidemic likely increased dramatically over time as infected hosts became transmissible. A single infected host sheds thousands of parasite spores (Troemel et al. 2008; Szumowski et al. 2012). A nonlinear relationship between dose and parasite acquisition, such as a threshold (McCallum et al. 2017), could collapse variation among hosts of different sizes: as parasite dose increases during an epidemic, all hosts regardless of size may ultimately contact enough parasites to become infected. Experiments in other host-parasite systems, like the Indian meal moth *Plodia interpunctella* and the bacterium *Bacillus thuringiensis* (Knell et al. 1996, 1998) or the spongy moth *Lymantria dispar* and nuclear polyhedrosis virus (Dwyer and Elkinton 1993), suggest that nonlinear transmission dynamics may be more common than previously thought. Newer theoretical work seeks to understand the emergent effects of nonlinearities on disease spread and epidemics (McCallum et al. 2017). We therefore suspect that in many host-parasite systems, the magnitude of transmission biases is dose dependent and should change as epidemics unfold.

Overall, we found evidence for our first hypothesis that small, young hosts are relatively unburdened by environmental parasites. Most species exhibit a similar positive correlation between age and body size (Randolph 1975; Weiner 2004; Cao et al. 2025). While the size difference between juvenile and adult *C. elegans* is quite large (roughly an order of magnitude; Byerly *et al*. 1976), superlinear relationships observed between body size and parasite burden in other systems suggest that even small size differences between juveniles and adults could generate substantial transmission biases (Mohr 1961; McCallum et al. 2017; Shaw et al. 2024). We expect that our results apply both to organisms with complex (multi-stage) life cycles and to those with simple (single-stage) life cycles. Although *C. elegans* does progress through a series of larval stages, body size varies continuously within each larval stage (Byerly *et al*. 1976), as is common in life cycles lacking discrete stages.

Given the relative health we found in small hosts, our second hypothesis was that natal dispersers formed during epidemics should be healthier than the populations they came from, since only small larvae (belonging to the L1 and L2 stages) can become dauer (dispersal) larvae. We found that dauers had lower odds of harboring *N. parisii* infections compared to the general population, and their infections were also less severe, especially in the more resistant CB4856 host strain (Figure 5). These results align with previous findings in fish (Poulin et al. 2012), arthropods (Altizer et al. 2000), and mammals (Folstad et al. 1991).

Disproportionately healthy dispersers have implications at both the individual and the metapopulation level. Parasite-free dispersers move faster and farther than infected dispersers (Bradley and Altizer 2005; Zilio et al. 2021) and can transition into reproductive adulthood more successfully once settled (Bubrig et al. 2022). We expect that size/age biases which protect young hosts can produce dispersers with higher-than-expected fitness and colonization success, thereby imposing selection to promote natal dispersal. If dispersers are preferentially healthy, we also expect reduced spread of parasites across a metapopulation. Reduced parasite load of dispersers should also delay parasite spread in the new habitat, enhancing the protection of juvenile hosts. Thus, the dynamics of transmission biases in one habitat may alter the trajectory of epidemics in other habitats through dispersal-mediated ecological feedbacks (Starrfelt and Kokko 2012).

In conclusion, we find that juvenile hosts are relatively shielded from infection compared to larger/older conspecifics, particularly at the start of epidemics. Juveniles may leverage this temporary advantage to disperse and avoid heavy infection, which we suspect could favor the evolution of parasite-mediated dispersal and shape how disease spreads among habitats. This study advances our understanding of the links between parasite spread and host dispersal and lays a foundation for future work disentangling how dispersal connects local and global spread of parasites.

## Acknowledgments

This work was supported by funding from the National Institute of General Medical Science (R35 GM137975-01). LTB was also supported by an NSF Research Training Grant (2021791).

## Conflict of Interest Statement

The authors declare no conflicts of interest.

## Author Contributions

LTB and IAA designed and performed assays, collected data, analyzed results, and wrote the manuscript. SKT and SGK performed assays, collected data, and analyzed results. AKG directed the study and edited the manuscript.

## Data Availability

Data and code can be accessed via Harvard Dataverse: (https://doi.org/10.7910/DVN/EEPMS8)

## Notes

### Competing Interest Statement

The authors have declared no competing interest.

