## Supplemental Figures/Tables for "Juvenile hosts and natal dispersers are protected in the early stages of epidemics"

**Supporting Information**

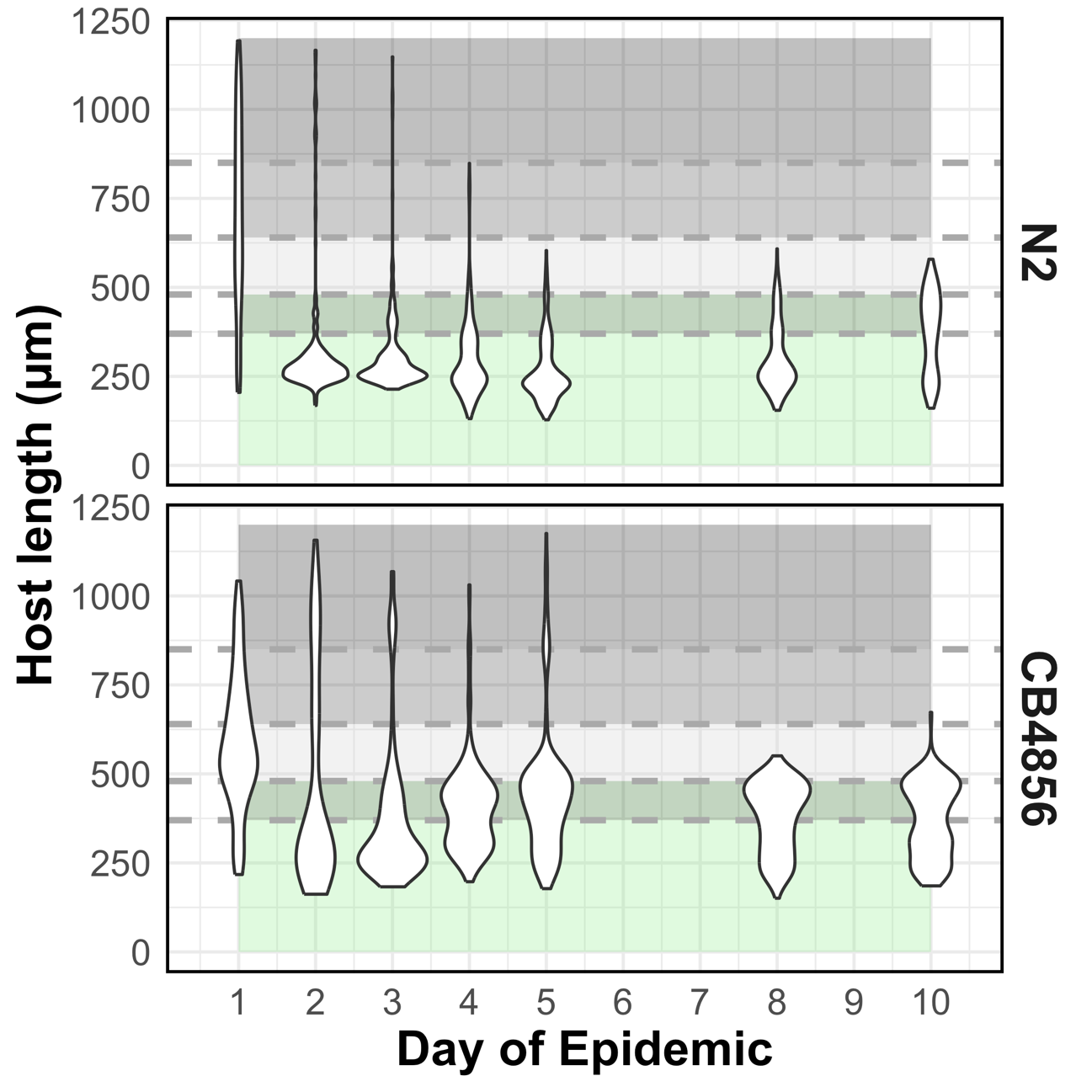

**Figure S1. Length in micrometers of *C. elegans* individuals in populations over time.** Length was measured from nose to tail (including curves of the body) in microscopy images. Horizontal dashed lines show the characteristic molting lengths between age classes, and shaded regions between lines correspond to the size ranges for each life stage. From bottom to top: 1) the L1-L2 molt at 370 µm, 2) the L2-L3 molt at 480 µm, 3) the L3-L4 molt at 640 µm, and 4) the L4-adult molt at 850 µm.

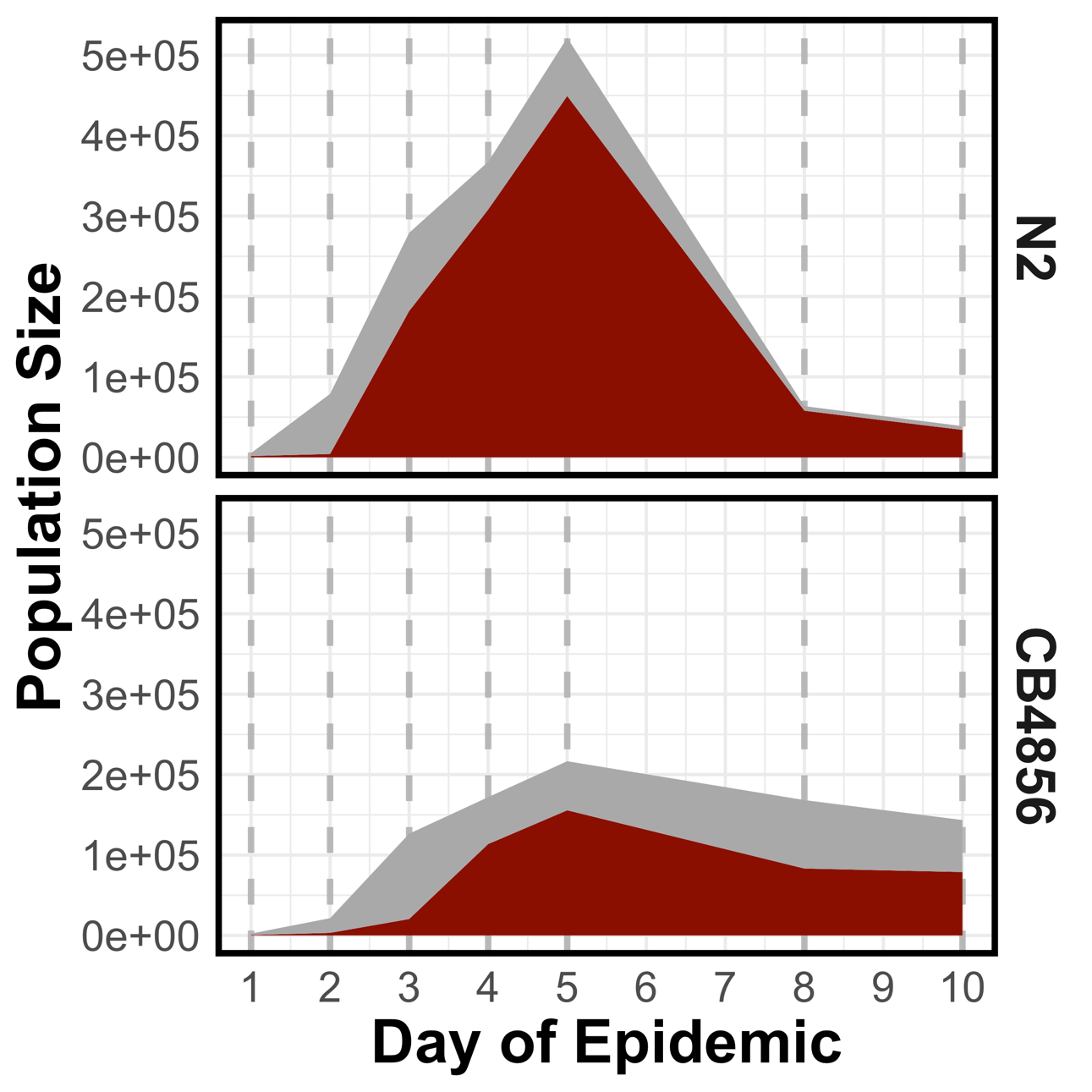

**Figure S2. Estimated population size and number of infected hosts over time.** The grey strip shows the estimated number of uninfected hosts while the red strip shows the estimated number of infected hosts. The number of uninfected versus infected hosts were estimated by multiplying each time point’s population size by its prevalence. Dashed lines indicate time points where these data were collected.

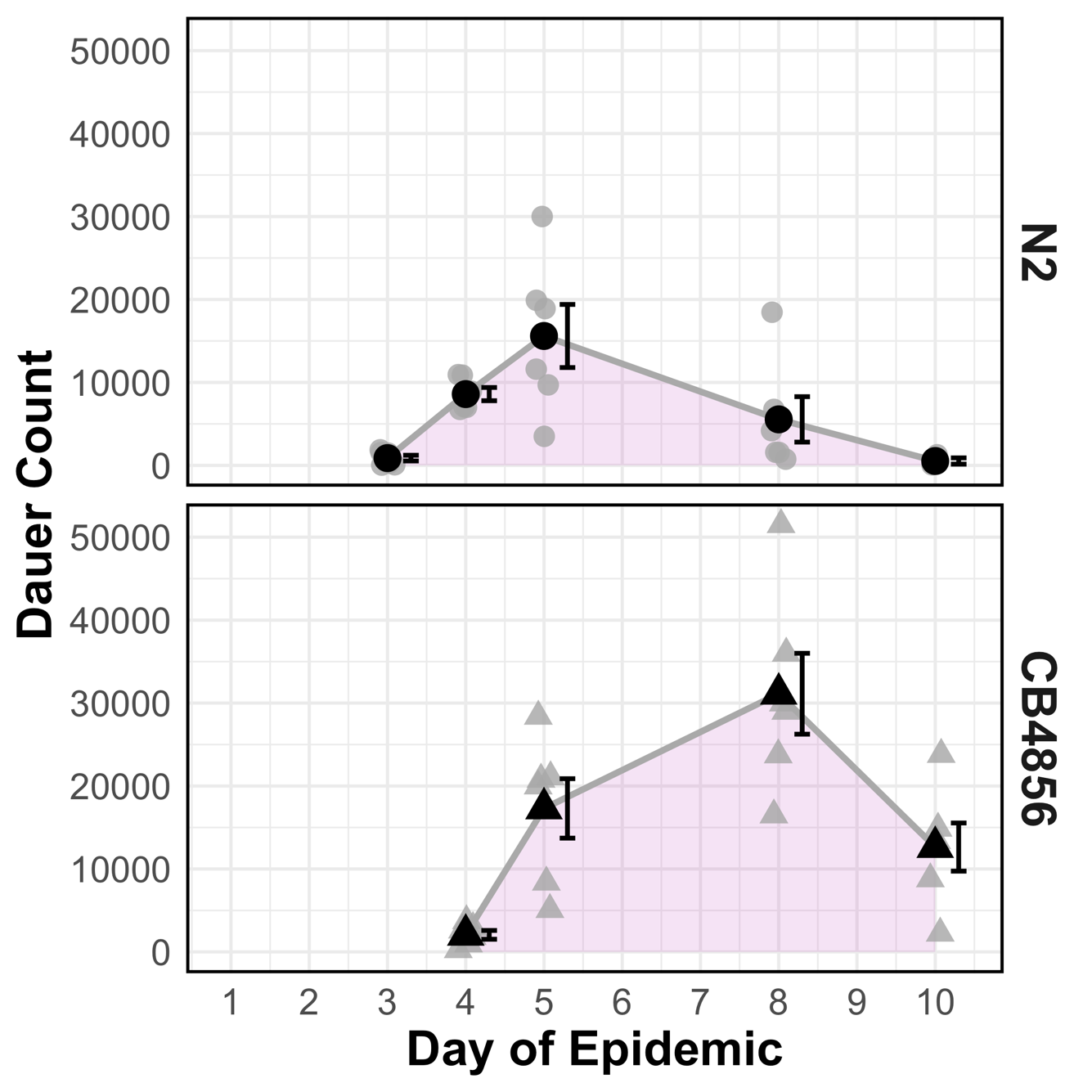

**Figure S3. Estimated size of the dauer pool over time.** Dauer pool counts were estimated from counting individuals in four replicate aliquots that survived a 15 minute 1% SDS treatment, which should include only dauers. Grey points show the estimated dauer count from a single plate, whereas black points show the mean of replicate plates. Error bars show the standard error of the mean.

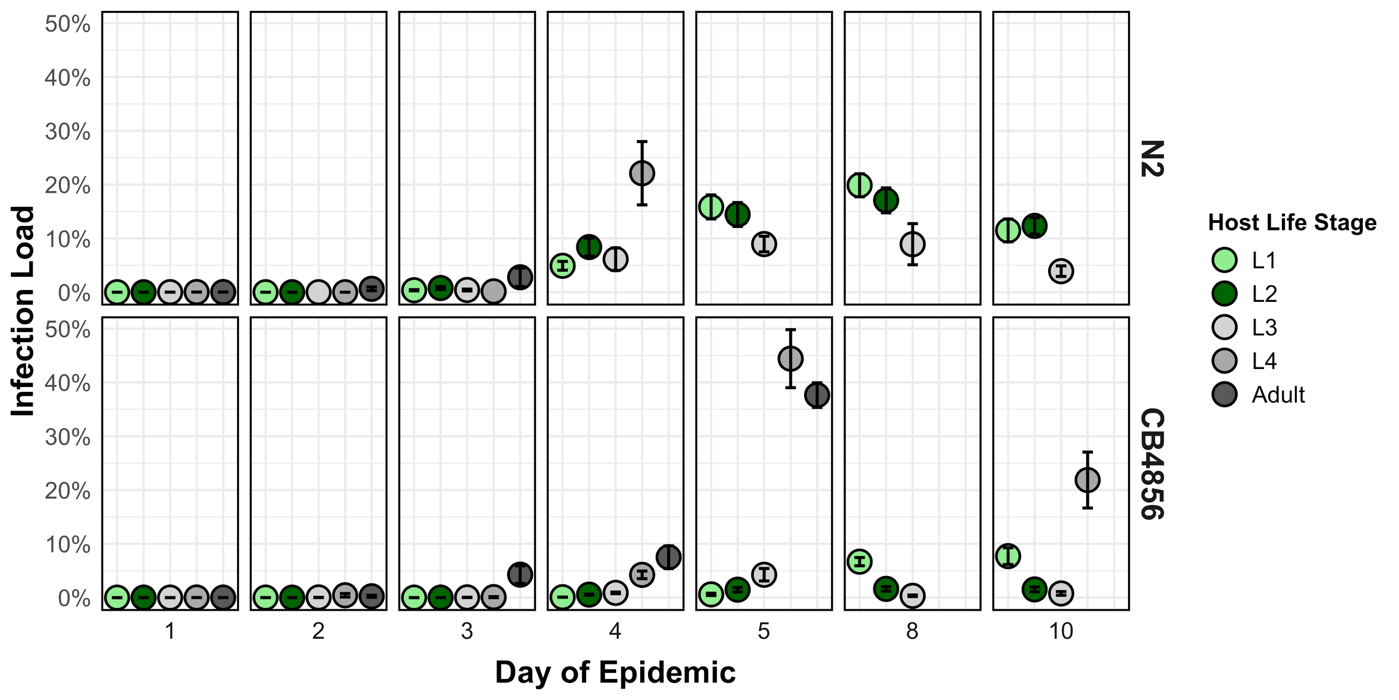

**Figure S4. The infection load of each life stage in the populations over time.** Each panel shows data from a single time point. Datapoints that are missing for particular life stages indicate that no members of that life stage were found at that time point. Error bars show standard error of the mean across replicate plates. Missing error bars indicate that the age class was only found on one of the replicate plates.

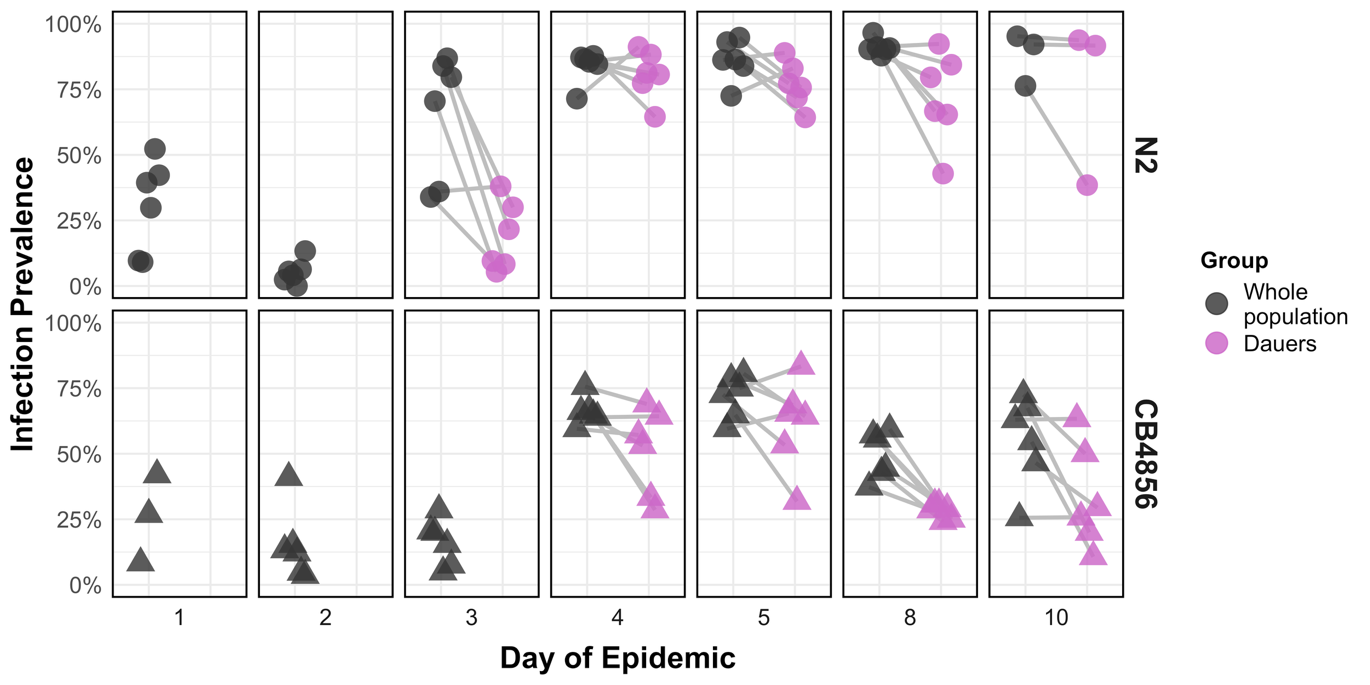

**Figure S5. Paired data showing the prevalence of dauer pools compared to their populations of origin.** Black points show the infection prevalence measured in the whole population (*i.e.,* including all life stages) for a given replicate plate. Pink points show the infection prevalence measured in the dauer pool for a given replicate plate. Lines connect datapoints that came from the same plate, to connect the prevalence measured in a whole population to the prevalence measured in that same population after being treated with 1% SDS to isolate the dauers.

| 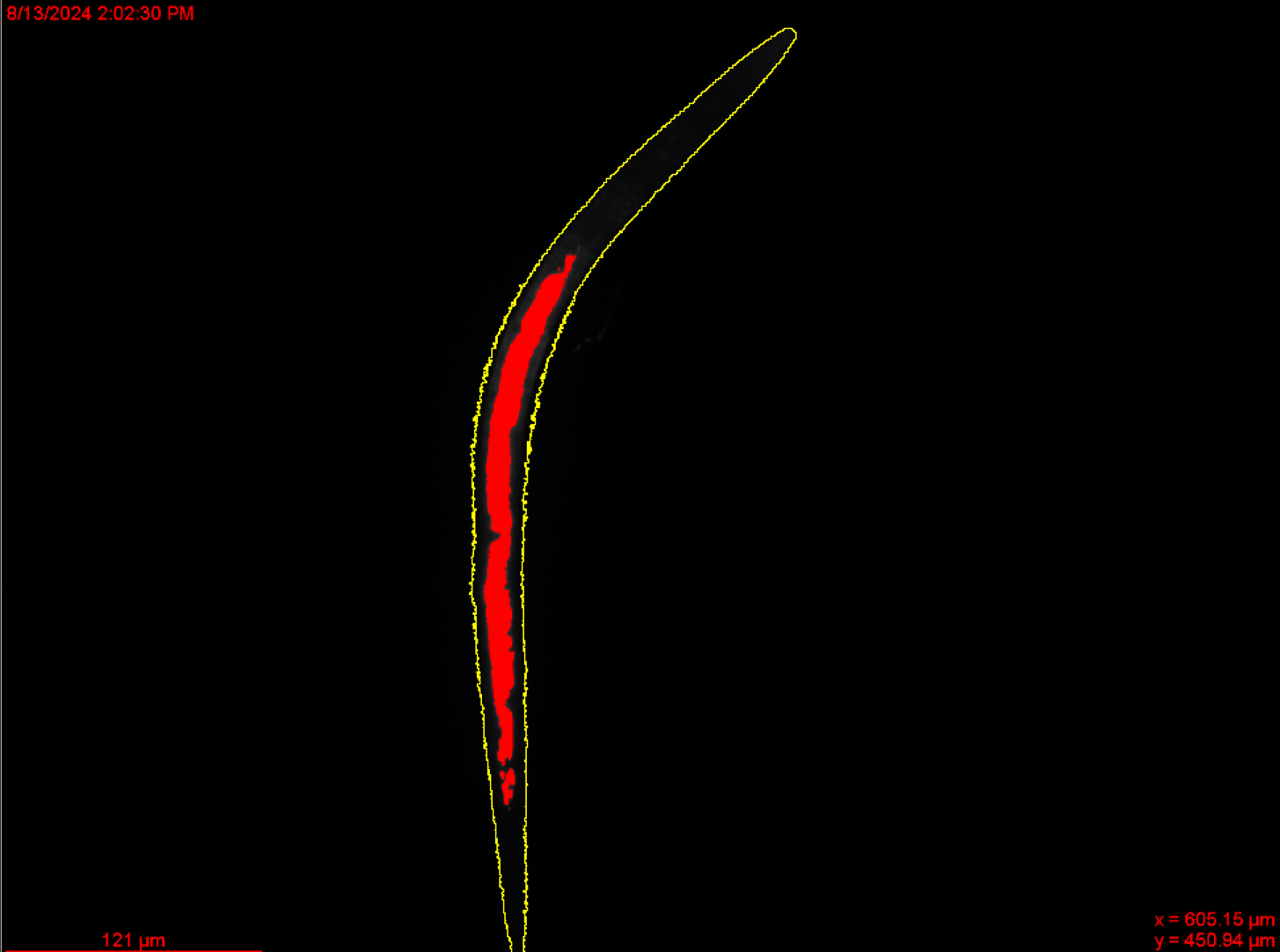 | 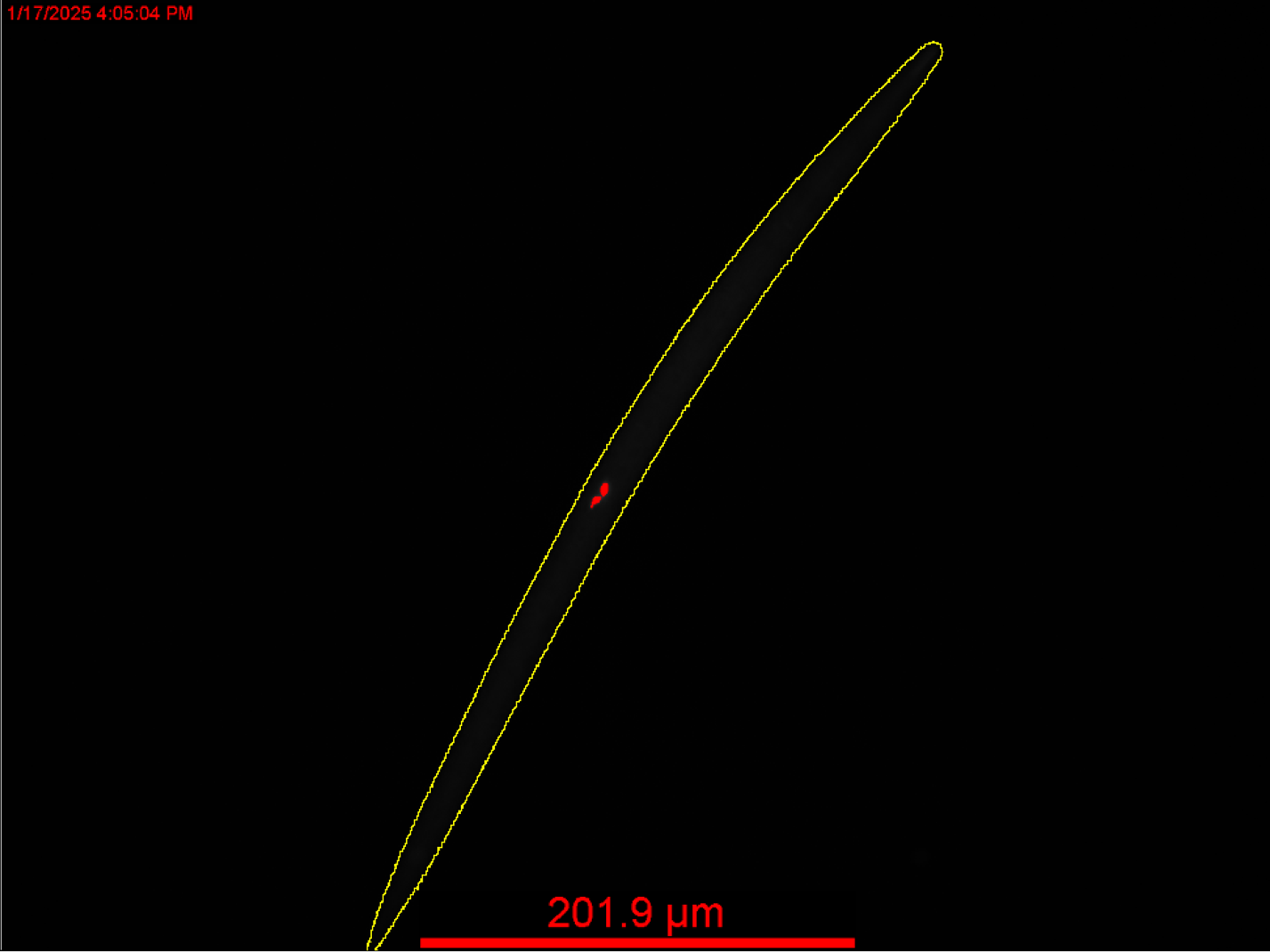 |
| --- | --- |
| **Figure S6. Representative FISH images of infected dauers.** Left: A typical infected N2 dauer on the last day of the epidemic (day 10). Right: A typical infected CB4856 on day 10. Nematodes are outlined in yellow. The red area inside of the nematodes indicates where *N. parisii* FISH probes bound. | |

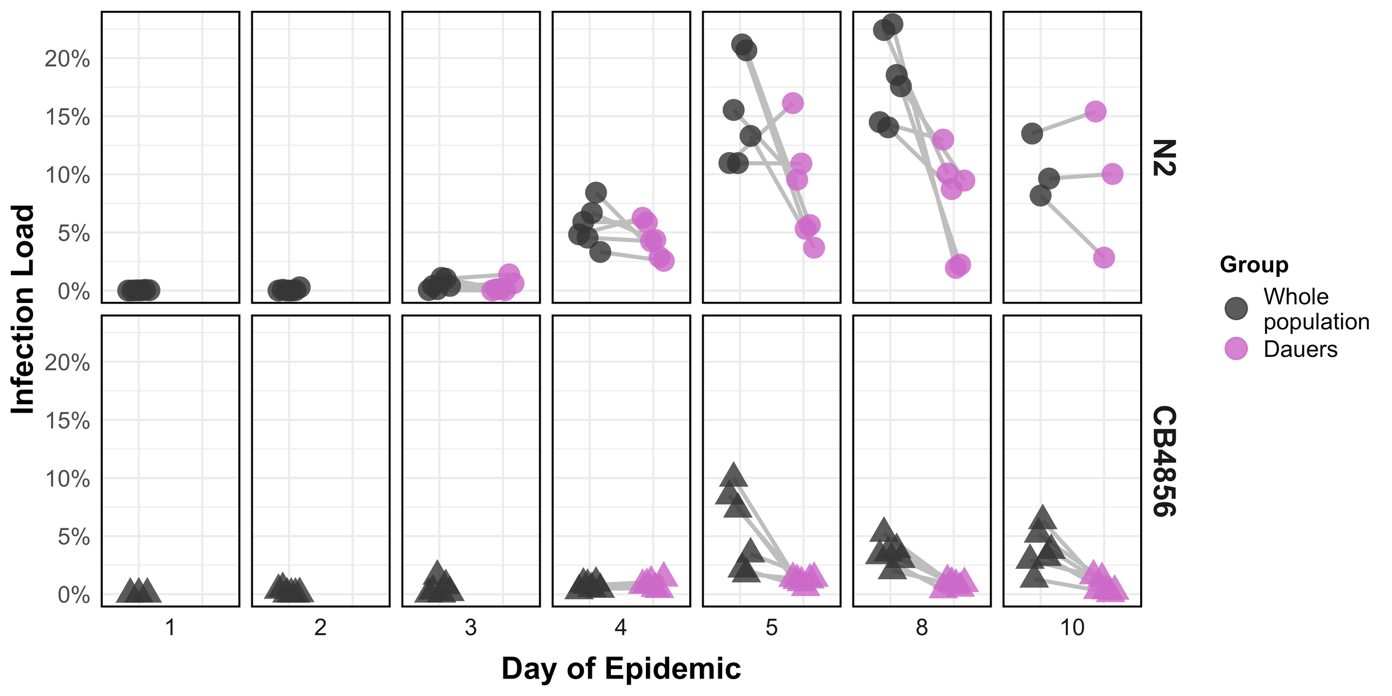

**Figure S7. Paired data showing the infection load of dauer pools compared to their populations of origin.** Black points show the infection load measured in the whole population (*i.e.,* including all life stages) for a given replicate plate. Pink points show the infection load measured in the dauer pool for a given replicate plate. Lines connect datapoints that came from the same plate, to connect the load measured in a whole population to the load measured in that same population after being treated with 1% SDS to isolate the dauers.

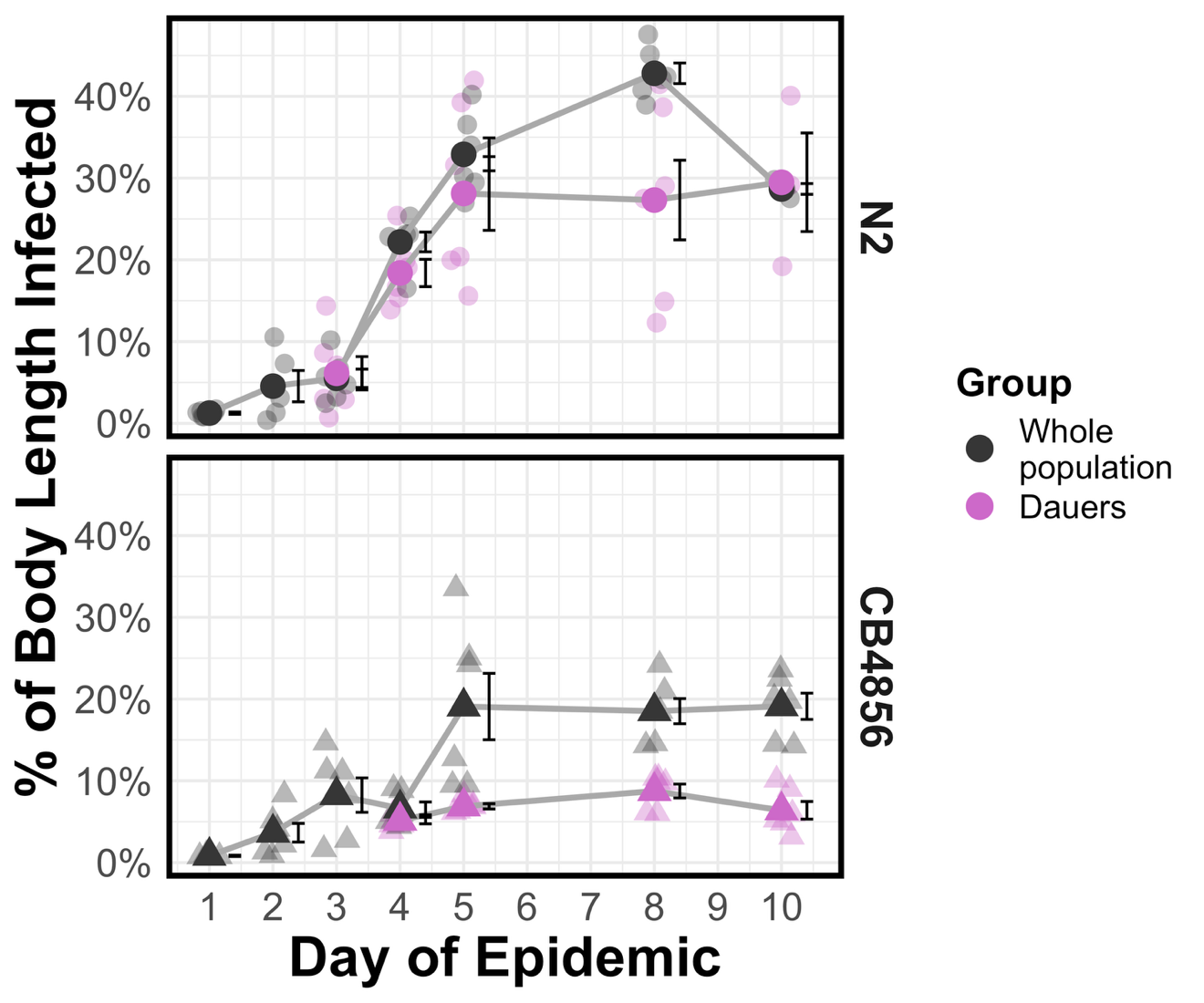

**Figure S8. Parasite linear load in the whole population versus the dauer pool.** Linear load was calculated from histograms of the fluorescent intensity along each host nematode’s head-to-tail axis. A host’s linear load is the percentage of that axis that has nonzero fluorescence, indicating parasite infection. Whole population data (black) may contain any *C. elegans* age classes including normal lifecycle larvae and dauers. Dauer pool data (pink) contain only individuals who survived SDS treatment, *i.e.* dauers. Days with no dauers in the population (see Figure S3) have no load data. Faded points show data from a single replicate population. Solid points show the mean across all replicate plates. Error bars show standard error of the mean.

**Table S1.** Typical lengths and timings of *C. elegans* life stages at 20°C, adapted from Byerly *et al.* 1976.

|  | **L1** | **L2** | **L3** | **L4** | **Adult** |
| --- | --- | --- | --- | --- | --- |
| **Length range** | < 370 µm | 370 - 480 µm | 480 - 640 µm | 640 - 850 µm | > 850 µm |
| **Time spent in stage** | 16 hours* | 8.5 hours | 9 hours | 12.5 hours | 1+ weeks |

*The L1-L2 molt occurs ~16 hours after hatching, but individuals arrest at the L1 stage if food is absent

**Table S2: Summary of analyses for predicting contact-rate with spores**

**A.** Model comparison

| **Model** | **Predictors** | **AIC** | ***χ*^2^** | ***p**** |
| --- | --- | --- | --- | --- |
| 1 | **LifeStage*Dose** + Block + (1\|ExposurePop) | 5057.8 | 1.446 | 0.971 |
| 2 | **LifeStage + Dose** + Block + (1\|ExposurePop) | 5055.2 |  |  |
| **Model (without interaction)** | **Predictors** | **AIC** | ***χ*^2^** | ***p**** |
| 1 | **LifeStage** + Dose + Block + (1\|ExposurePop) | 5055.2 | 62.666 | <0.0001 |
| 2 | Dose + Block + (1\|ExposurePop) | 5113.9 |  |  |

Each row corresponds to a different statistical model. Columns show the model terms, the AIC scores, the chi-square values, and the *Bonferroni corrected *p values*. Bold model terms show the variable of interest.

**B.** Summary of winning model

*Model: SporeCount ~ LifeStage + Dose + Block + (1|ExposurePop)*

*GLMM, negative binomial distribution, log link*

*Ref: L1 LifeStage, Block 1*

| **Predictor** | **Level** | **Coefficient ± SE** | | | **Multiplicative change in spore count (95% CI)** | | **z-value** |
| --- | --- | --- | --- | --- | --- | --- | --- |
| (Intercept) |  | | 1.837 ± 0.259 | 6.277 (4.842, 8.136) | | 7.079 | |
| LifeStage | L4 | | 1.271 ± 0.173 | 3.565 (2.999, 4.239) | | 7.349 | |
|  | Adult | | 1.971 ± 0.167 | 7.180 (6.073, 8.489) | | 11.771 | |
| Dose |  | | 0.013 ± 0.002 | 1.013 (1.011, 1.016) | | 5.367 | |
| Block | 2 | | -0.487 ± 0.219 | 0.615 (0.494, 0.765) | | -2.225 | |
|  | 3 | | -0.204 ± 0.191 | 0.815 (0.674, 0.987) | | -1.069 | |

**Table S3: Summary of analyses for predicting infection status of hosts**

**A.** Model comparison

| **Model** | **Predictors** | **AIC** | ***χ*^2^** | ***p*** |
| --- | --- | --- | --- | --- |
| 1 | **DayOfEpidemic*RescaledLength** + Strain+ Block + (1\|UniquePlateID) | 3392.1 | 63.35 | < 0.0001 |
| 2 | **DayOfEpidemic + RescaledLength** + Strain+ Block + (1\|UniquePlateID) | 3453.4 |  |  |

Each row corresponds to a different statistical model. Columns show the model terms, the AIC scores, chi-square values, and *p values*. Bold model terms show the variable of interest. For model fitting and interpretation, the original lengths (in units of µm) were divided by 100. Therefore, model coefficients for length should be interpreted as the change per 100 µm of length

**B.** Summary of winning model

*Model: IsInfected ~ DayOfEpidemic*RescaledLength + Strain + Block + (1|ExposurePop)*

*GLMM, binomial distribution, logit link*

*Ref: N2 Strain, Block 1*

| **Predictor** | **Level** | **Coefficient ± SE** | | | **Odds Ratio (95% CI)** | | **z-value** |
| --- | --- | --- | --- | --- | --- | --- | --- |
| (Intercept) |  | | -5.169 ± 0.562 | 0.006 (0.003, 0.010) | | -9.189 | |
| DayOfEpidemic |  | | 0.918 ± 0.093 | 2.505 (2.282, 2.749) | | 9.867 | |
| RescaledLength |  | | 0.811 ± 0.058 | 2.250 (2.123, 2.385) | | 13.931 | |
| DayOfEpidemic: RescaledLength |  | | -0.098 ± 0.013 | 0.907 (0.895, 0.918) | | -7.791 | |
| Strain | CB4856 | | -1.519 ± 0.413 | 0.219 (0.145, 0.331) | | -3.675 | |
| Block | 2 | | 0.095 ± 0.411 | 1.100 (0.729, 1.659) | | 0.232 | |

**Table S4: Summary of analyses for predicting the parasite load of infected hosts**

**A.** Model comparison

| **Model** | **Predictors** | **AIC** | ***χ*^2^** | ***p**** |
| --- | --- | --- | --- | --- |
| 1 | **DayOfEpidemic*RescaledLength** + Strain + Block + (1\|UniquePlateID) | -8228.8 | 3.342 | 0.1350 |
| 2 | **DayOfEpidemic + RescaledLength** + Strain + Block + (1\|UniquePlateID) | -8227.4 |  |  |
| **Model (without interaction)** | **Predictors** | **AIC** | ***χ*^2^** | ***p**** |
| 1 | DayOfEpidemic + **RescaledLength** + Strain + Block + (1\|UniquePlateID) | -8227.4 | 36.509 | <0.0001 |
| 2 | DayOfEpidemic + Strain + Block + (1\|UniquePlateID) | -8192.9 |  |  |

Each row corresponds to a different statistical model. Columns show the model terms, the AIC scores, the chi-square values, and the *Bonferroni corrected *p values*. Bold model terms show the variable of interest.

**B.** Summary of winning model

*Model: LoadPercent ~ DayOfEpidemic + RescaledLength + Block + (1|UniquePlateID)*

*GLMM, beta distribution, logit link, zero inflated formula: ~1*

*Ref: N2 Strain, Block 1*

| **Predictor** | **Level** | **Coefficient ± SE** | | | **Multiplicative change in load (95% CI)** | | **z value** |
| --- | --- | --- | --- | --- | --- | --- | --- |
| (Intercept) |  | | -4.441 ± 0.214 |  | | -20.767 | |
| DayOfEpidemic |  | | 0.281 ± 0.028 | 1.325 (1.288, 1.362) | | 10.056 | |
| RescaledLength |  | | 0.106 ± 0.017 | 1.112 (1.093, 1.131) | | 6.171 | |
| Strain | CB4856 | | -0.822 ± 0.150 | 0.440 (0.379, 0.511) | | -5.495 | |
| Block | 2 | | 0.013 ± 0.149 | 1.013 (0.873, 1.175) | | 0.086 | |

**Table S5: Summary of analyses for predicting the parasite prevalence of the whole population versus dauer pool**

**A.** Model comparison

| **Model** | **Predictors** | **AIC** | ***χ*^2^** | ***p*** |
| --- | --- | --- | --- | --- |
| 1 | **SDS** + DayOfEpidemic + Strain + Block + (1\|UniquePlateID) | 5655.7 | 143.41 | <0.0001 |
| 2 | DayOfEpidemic + Strain + Block + (1\|UniquePlateID) | 5797.1 |  |  |

Each row corresponds to a different statistical model. Columns show the model terms, the AIC scores, the chi-square values, and the associated *p values*. Bold model terms show the variable of interest.

**B.** Summary of winning model

*Model: IsInfected ~ SDS + DayOfEpidemic + Strain + Block + (1|UniquePlateID)*

*GLMM, binomial distribution, logit link*

*Ref: Pre SDS, N2 Strain, Block 1*

| **Predictor** | **Level** | **Coefficient ± SE** | | | **Odds Ratio (95% CI)** | **z value** |
| --- | --- | --- | --- | --- | --- | --- |
| (Intercept) |  | | -0.901 ± 0.350 | 0.406 (0.286, 0.577) | | -2.573 |
| SDS | Post | | -0.906 ± 0.077 | 0.404 (0.374, 0.436) | | -11.806 |
| DayOfEpidemic |  | | 0.364 ± 0.055 | 1.440 (1.362, 1.522) | | 6.568 |
| Strain | CB4856 | | -1.182 ± 0.314 | 0.307 (0.224, 0.419) | | -3.771 |
| Block | 2 | | -0.194 ± 0.313 | 0.824 (0.603, 1.126) | | -0.620 |

**Table S6: Summary of analyses for predicting the parasite load of the whole population versus dauer pool**

**A.** Model comparison

| **Model** | **Predictors** | **AIC** | ***χ*^2^** | ***p**** |
| --- | --- | --- | --- | --- |
| 1 | **Strain*SDS** + DayOfEpidemic + Block + (1\|UniquePlateID) | -11790 | 21.579 | <0.0001 |
| 2 | **Strain + SDS** + DayOfEpidemic + Block + (1\|UniquePlateID) | -11770 |  |  |
| **Model (N2 only)** | **Predictors** | **AIC** | ***χ*^2^** | ***p**** |
| 1 | **SDS** + DayOfEpidemic + Block + (1\|UniquePlateID) | -6621.8 | 87.571 | <0.0001 |
| 2 | DayOfEpidemic + Block + (1\|UniquePlateID) | -6538.2 |  |  |
| **Model (CB4856 only)** | **Predictors** | **AIC** | ***χ*^2^** | ***p**** |
| 1 | **SDS** + DayOfEpidemic + Block + (1\|UniquePlateID) | -5229.7 | 114.47 | <0.0001 |
| 2 | DayOfEpidemic + Block + (1\|UniquePlateID) | -5117.3 |  |  |

Each row corresponds to a different statistical model. Columns show the model terms, the AIC scores, the chi-square values, and the *Bonferroni corrected *p values*. Bold model terms show the variable of interest.

**B.** Summary of winning N2 model

*Model: LoadPercent ~ SDS + DayOfEpidemic + Block + (1|UniquePlateID)*

*GLMM, beta distribution, logit link, dispersion formula: ~SDS + DayOfEpidemic*

*Ref: Pre SDS, Block 1*

| **Predictor** | **Level** | **Coefficient ± SE** | | | **Multiplicative change in load (95% CI)** | | **z value** |
| --- | --- | --- | --- | --- | --- | --- | --- |
| (Intercept) |  | | -4.641 ± 0.237 |  | | -19.588 | |
| SDS | Post | | -0.338 ± 0.042 | 0.713 (0.684, 0.744) | | -8.066 | |
| DayOfEpidemic |  | | 0.399 ± 0.040 | 1.490 (1.431, 1.552) | | 9.884 | |
| Block | 2 | | -0.121 ± 0.212 | 0.886 (0.717, 1.095) | | -0.572 | |

**C.** Summary of winning CB4856 model

*Model: LoadPercent ~ SDS + DayOfEpidemic + Block + (1|UniquePlateID)*

*GLMM, beta distribution, logit link, dispersion formula: ~SDS + DayOfEpidemic*

*Ref: Pre SDS, Block 1*

| **Predictor** | **Level** | **Coefficient ± SE** | | | **Multiplicative change in load (95% CI)** | | **z value** |
| --- | --- | --- | --- | --- | --- | --- | --- |
| (Intercept) |  | | -3.930 ± 0.170 |  | | -23.087 | |
| SDS | Post | | -1.020 ± 0.087 | 0.361 (0.331, 0.393) | | -11.752 | |
| DayOfEpidemic |  | | 0.150 ± 0.024 | 1.162 (1.134, 1.190) | | 6.238 | |
| Block | 2 | | 0.031 ± 0.096 | 1.031 (0.937, 1.135) | | 0.318 | |

**Table S7: Summary of analyses for predicting the linear load of the whole population versus dauer pool**

**A.** Model comparison

| **Model** | **Predictors** | **AIC** | ***χ*^2^** | ***p**** |
| --- | --- | --- | --- | --- |
| 1 | **Strain*SDS** + DayOfEpidemic + Block + (1\|UniquePlateID) | -5004.6 | 19.346 | <0.0001 |
| 2 | **Strain + SDS** + DayOfEpidemic + Block + (1\|UniquePlateID) | -4987.2 |  |  |
| **Model (N2 only)** | **Predictors** | **AIC** | ***χ*^2^** | ***p**** |
| 1 | **SDS** + DayOfEpidemic + Block + (1\|UniquePlateID) | -2831.3 | 18.172 | <0.0001 |
| 2 | DayOfEpidemic + Block + (1\|UniquePlateID) | -2815.2 |  |  |
| **Model (CB4856 only)** | **Predictors** | **AIC** | ***χ*^2^** | ***p**** |
| 1 | **SDS** + DayOfEpidemic + Block + (1\|UniquePlateID) | -2238.2 | 35.768 | <0.0001 |
| 2 | DayOfEpidemic + Block + (1\|UniquePlateID) | -2204.4 |  |  |

Each row corresponds to a different statistical model. Columns show the model terms, the AIC scores, the chi-square values, and the *Bonferroni corrected *p values*. Bold model terms show the variable of interest.

**B.** Summary of winning N2 model

*Model: LinearLoad ~ SDS + DayOfEpidemic + Block + (1|UniquePlateID)*

*GLMM, beta distribution, logit link, dispersion formula: ~1*

*Ref: Pre SDS, Block 1*

| **Predictor** | **Level** | **Coefficient ± SE** | | | **Multiplicative change in load (95% CI)** | | **z value** |
| --- | --- | --- | --- | --- | --- | --- | --- |
| (Intercept) |  | | -2.852 ± 0.188 |  | | -15.191 | |
| SDS | Post | | -0.162 ± 0.038 | 0.850 (0.818, 0.883) | | -4.255 | |
| DayOfEpidemic |  | | 0.300 ± 0.032 | 1.349 (1.307, 1.393) | | 9.336 | |
| Block | 2 | | -0.149 ± 0.174 | 0.862 (0.724, 1.026) | | -0.855 | |

**C.** Summary of winning CB4856 model

*Model: LinearLoad ~ SDS + DayOfEpidemic + Block + (1|UniquePlateID)*

*GLMM, beta distribution, logit link, dispersion formula: ~1*

*Ref: Pre SDS, Block 1*

| **Predictor** | **Level** | **Coefficient ± SE** | | | **Multiplicative change in load (95% CI)** | | **z value** |
| --- | --- | --- | --- | --- | --- | --- | --- |
| (Intercept) |  | | -2.485 ± 0.114 |  | | -21.781 | |
| SDS | Post | | -0.405 ± 0.069 | 0.667 (0.623, 0.715) | | -5.867 | |
| DayOfEpidemic |  | | 0.115 ± 0.016 | 1.121 (1.103, 1.140) | | 7.039 | |
| Block | 2 | | -0.050 ± 0.089 | 0.951 (0.870, 1.039) | | -0.567 | |
